# Economic Burden of Hypoglycaemia among Type II Diabetes Mellitus Patients in Malaysia

**DOI:** 10.1101/518225

**Authors:** Syed Mohamed Aljunid, Yin New Aung, Aniza Ismail, Siti Athirah Zafirah Abdul Rashid, Amrizal M Nur, Julius Cheah, Priya Matzen

## Abstract

**Objective:** The main aim of this study is to identify the direct cost and economic burden of hypoglycaemia among patients with Type II diabetes mellitus in Malaysia.

**Methods:** The incurred cost for hypoglycaemia among patients admitted to University Kebangsaan Malaysia Medical Centre (UKMMC) was explored from a cross sectional study. 20-79 year patients discharged between Jan 2010 to Sept 2015 and having an ICD-10 code of hypoglycaemia as a primary diagnosis in the casemix database were included in the study. Costing analysis from the perspective of health providers was completed using step-down approach. Data related to hospital cost were collected using hospital-costing template, based on three levels of cost centres. The costing data from UKMMC was used to estimate the national burden of hypoglycaemia among type II diabetics for the whole country.

**Results:** Of 244 diabetes patients admitted primarily for hypoglycaemia to UKMMC, 52% were female and 88% were over 50 years old. The cost increased with severity. Managing a hypoglycaemic case requires 5 days (median) of inpatient stay with a range of 2-26 days; and costs RM 9,083 (USD 2,323). 30% of the cost came from the ward services cost (final cost centre), 16% of the cost came from ICU services and 15% from pharmacy services(secondarylevel cost centres). Based on the prevalence of hypoglycaemia-related admissions of 3.2%, the total cost of care for hypoglycaemia among adult diabetes in Malaysia is estimated to be RM 117.4 (USD 30.0) million, which is translated as 0.5% of Ministry of Health budget.

**Conclusion:** Hypoglycaemia imposes substantial economic impact even without the direct and indirect cost incurred by patients and other cost of complications. Proper diabetic care and health education is needed in diabetic management to reduce episodes of hypoglycaemia.

## Introduction

Hypoglycaemia is a not an uncommon presentation at accident and emergency (A & E) department. An estimate of 2-4% of diabetes deaths is due to hypoglycaemia [1]. Hypoglycaemic episode is inevitable in 90% of diabetics with insulin therapy[2], and those with certain antihyperglycaemic medicine are also at risk. Elderly and those with comorbid conditions are more likely to get severe hypoglycaemia. [3] The incidence of hypoglycaemia varies with studies. On average, type 1 diabetics had two symptomatic hypoglycaemia episodes per week [4]and one severe hypoglycaemia episode per year[5]; with a lesser occurrence among type II diabetes patients.[2] However, type II diabetes with progressive insulin deficiency, long duration of diabetes, and tight glycaemic control also have similar risk of hypoglycaemia as type 1 diabetes.[5]

Eleven percent of type 2 diabetic patients with different anti-diabetic regimens suffered one or more hypoglycaemic episodes per year. Miller et al reported that 24.5% of patients reported at least one hypoglycaemic episode in 3 months period [6, 7]. Mild hypoglycaemic attacks occurs frequently, as high as 1-2 episodes per week[8] and severe hypoglycaemic attacks occurs less frequently with an incidence rate of 1 to 2.7 episodes by per patient per year [8, 9].

DM is one of the commonest chronic non-communicable diseases globally. Its incidence and prevalence are escalating, and Asia-Pacific region is at the forefront of the current epidemic. As of 2015, it is estimated that 8.8% of adult population in Southeast Asia region has diabetes [10]. Prevalence of DM among Malaysian over 20-79 years is estimated to be 17.5%[11], ranking third in Asia-Pacific region. [12, 13] It was estimated that in 2030, Malaysia would have a total number of 2.48 million diabetics compared to 0.94 million in 2000 that can be translated as 164% increase [14].

The chronic nature of diabetes and its devastating complications make it a very costly disease. To prevent the risk of acute and chronic complication, diabetic patients required continuous medical care [2, 15]. Hypoglycaemia is unpredictable and undesirable side effects among diabetic patients. Frequent and potentially fatal complication may occur among hypoglycaemic patients with Type 1 or Type 2 diabetes treated with insulin, and in patients with Type 2 diabetes treated with certain oral anti-diabetic medicine.

Hypoglycaemic attacks require defence mechanism against falling serum glucose, and frequent attack result in increasing cycles of recurrent hypoglycaemia [13]. Fear of hypoglycaemia is not uncommon in patients with diabetes. Experiences of severe hypoglycaemic episodes increase the fear for future hypoglycaemic event. Approximately 40% of patients admitted that fear of hypoglycaemia caused them to maintain their blood glucose levels at higher than recommended values [14]. This questions compliance of anti-diabetic medicine with possible complication due to uncontrolled diabetes and increased health care cost. Regardless of bearing a significant burden by hypoglycaemia, the cost of hypoglycaemia in Malaysia is yet known. This study identifies the direct cost and economic burden of hypoglycaemia among patients with type II diabetes mellitus on insulin in Malaysia.

## Methods

A cross sectional study was conducted to identify the incurred cost due to hypoglycaemia among patients admitted to Universiti Kebangsaan Malaysia Medical Centre (UKMMC). Using the electronic medical database kept at UKMMC, patients discharged from January 2010 to September 2015 were classified into diagnosis related groups (DRG) with MY-DRG^®^ grouping software. 20-79 year old patients having hypoglycaemia as a primary reason for admission in the casemix database is identified using ICD 10 codes. The ICD 10 codes associated with hypoglycaemia: E16.0 (Drug Induced Hypoglycaemia without coma, E16.1 (Other Hypoglycaemia) and E16.2 (Hypoglycaemia Unspecified) were included in the study. Costing analysis was carried out using step-down approach. Financial staffs in the hospital were given a costing template to retrieve financial data from the hospital records. The templates classifies the departments into three levels of cost centres: overhead cost centres (e.g.; administration, utilities, maintenance etc.), intermediate cost centres (e.g.; pharmacy, radiology etc.) and final cost centres (all wards and all clinics). Information on financial expenditures and output for each cost centre was recorded. The information recoded includes the total expenditure, total number discharges, inpatient days, number of patient visits for outpatient clinics and floor space. All the information of the activities reflecting the workload such as number of discharges, inpatient days, floor space and number of outpatient visits were gathered for appropriate allocation.

Both capital cost (building, equipment and furniture cost) and recurrent cost (staff salary and other recurrent cost) were combined in estimating the cost for each cost centre. The final allocated costs for each inpatient cost centreswere then divided by the total units of inpatient days to obtain the cost of providing services on a per-patient per-day of stay basis, which is referred as unit cost. The unit cost is finally multiplied with the individual patient’s length of stay to obtain the cost of care per patient per discharge. All these steps were simplified by using the Clinical Cost Modelling Software Version 2.1 (CCM Ver. 2.1). CCM is the step-down costing tool that is being used by the casemix system in UKMMC.

The cost is triangulated by developing clinical pathways for a hypoglycaemic episode. An expert group meeting was held to develop the common clinical pathways of managing hypoglycaemia. These experts include physicians, endocrinologists, and pharmacists who are working in the A&E department and also in the clinical departments of the hospital.

In imputing the national burden, estimates for incidence and prevalence of diabetes were obtained from NMHS 2015 [8] and the International Diabetes Federation.[9] The possible episode of hypoglycaemia incidence was estimated from the Malaysian Hypoglycaemia Assessment Tool Study (HAT study) [16]. The unit cost for management of hypoglycaemia calculated in the step-down approach was used to estimate the burden. Sensitivity analysis was carried out by varying the incidence of hypoglycaemia to obtain the worst-case scenario and best-case scenario.

## Results

903 cases (0.54% of the total cases) discharged from UKMMC between January 2010 and September 2015 had a diagnosis of hypoglycaemia (Table 1).

**Table 1.** Number of Hypoglycaemia Patients By Year in UKMMC Casemix Database

| Year | Nos. | Total Discharges | % Hypoglycaemia of Total Discharges |
| --- | --- | --- | --- |
| 2010 | 199 | 32,144 | 0.62% |
| 2011 | 157 | 26,262 | 0.60% |
| 2012 | 180 | 28,728 | 0.63% |
| 2013 | 177 | 33,473 | 0.53% |
| 2014 | 119 | 27,303 | 0.44% |
| 2015 | 71 | 18,623 | 0.38% |
| **Total** | 903 | 166,533 | 0.54% |

Among these cases, 33.4% were classified into MY-DRG^®^ casemix groups of Endocrine, Nutrition and Metabolism Group followed by Respiratory System Group (12.6%) and Cardiovascular System Group (9.7%). Central Nervous System and Nephro-urinary System contributed 6.4% of the hypoglycaemia individually (Table 2).

**Table 2.** Hypoglycemia by Casemix Main Group (CMG)

| No | Casemix Main Groups (CMG) | Frequency | Percent |
| --- | --- | --- | --- |
| 1 | Endocrine system, nutrition & metabolism Groups | 302 | 33.4 |
| 2 | Respiratory system Groups | 114 | 12.6 |
| 3 | Cardiovascular system Groups | 88 | 9.7 |
| 4 | Central nervous system Groups | 58 | 6.4 |
| 5 | Nephro-urinary System Groups | 58 | 6.4 |
| 6 | Infectious & parasitic diseases Groups | 51 | 5.6 |
| 7 | Musculoskeletal system & connective tissue Groups | 47 | 5.2 |
| 8 | Digestive system Groups | 45 | 5 |
| 9 | Hepatobiliary & pancreatic system Groups | 38 | 4.2 |
| 10 | Female reproductive system Groups | 26 | 2.9 |
| 11 | Skin, subcutaneous tissue & breast Groups | 23 | 2.5 |
| 12 | Haemopoietic& immune system Groups | 17 | 1.9 |
| 13 | Ear, nose, mouth & throat Groups | 10 | 1.1 |
| 14 | Myeloproliferative system & neoplasms Groups | 7 | 0.8 |
| 15 | Mental Health and Behavioral Groups | 5 | 0.6 |
| 16 | Injuries, poisonings & toxic effects of drugs Groups | 4 | 0.4 |
| 17 | Eye and Adnexa Groups | 3 | 0.3 |
| 18 | Male reproductive System Groups | 2 | 0.2 |
| 19 | Deliveries Groups | 2 | 0.2 |
| 20 | Factors influencing health status & other contacts with health services Groups | 2 | 0.2 |
| 21 | Substance abuse & dependence Groups | 1 | 0.1 |
| Total |  | 903 | 100 |

Most of the hypoglycaemic conditions were admitted to the medical ward (91.7%), and 8.1% were admitted to the surgical unit.

Of the 302 hypoglycaemic cases with endocrine problems, diabetes in particular, 244 cases were admitted primarily for hypoglycaemia and 58 were admitted for hypoglycaemia as a secondary reason (Table 3).

**Table 3.**
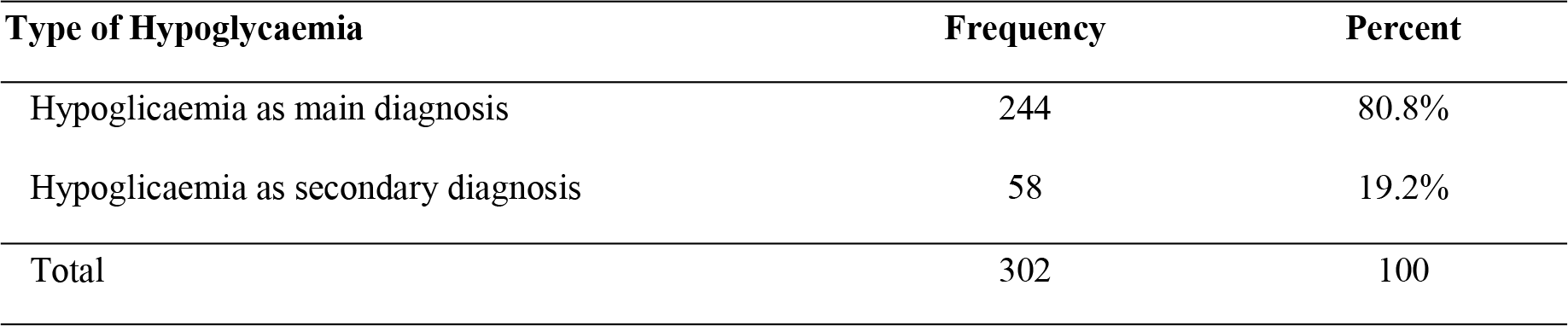
Type of Hypoglycemia among cases in CMG Endocrine System, Nutrition & Metabolism

| Type of Hypoglycaemia | Frequency | Percent |
| --- | --- | --- |
| Hypoglycaemia as main diagnosis | 244 | 80.8% |
| Hypoglycaemia as secondary diagnosis | 58 | 19.2% |
| Total | 302 | 100 |

Only 244 cases with hypoglycaemia as primary reason were included in the study for further analysis. Females were more likely to suffer from hypoglycaemia, as almost 52% of the cases were female. 99.2% of all cases were discharged home well. In the MY-DRG^®^ system, patients are classified into one of the three levels of severity. Cases in severity level 1 do not have complications or co-morbidities where as those in severity level 2 have minor complications and co-morbidities. Patients in severity level 3 have major complications and co-morbidities. About 57.4% of the primary hypoglycaemia cases were in severity level 3. Most (88.1%) of the patients suffering from hypoglycaemic attacks were in the age group of 50-79 years old, with 55.3% of the cases required not more than 5 days of hospital admission (Table 4).

**Table 4:** Characteristics of hypoglycemic cases among CMG Endocrine System, Nutrition and Metabolism

| Characteristics |  | Hypoglycemia as primary diagnosis<br>(n =244) |
| --- | --- | --- |
| Gender (Female, %) |  | 127 (52%) |
| Discharged status | - Discharged to home | 242 (99.2%) |
|  | - Others | 2 (0.8%) |
| Severity Level | - Severity Level 1 | 23 (9.4%) |
|  | - Severity Level 2 | 81(33.2%) |
|  | - Severity Level 3 | 140 (57.4%) |
| Age group | - 20 - 29 years | 3 (1.2%) |
|  | - 30 - 39 years | 8 (3.3%) |
|  | - 40 - 49 years | 18 (7.4%) |
|  | - 50 - 59 years | 34 (13.9%) |
|  | - 60 - 69 years | 84 (34.4%) |
|  | - 70 - 79 years | 97 (39.8%) |
| Length of stay | - <=5 days | 135 (55.3%) |
|  | - 6 – 10 days | 77 (31.6%) |
|  | - 11 – 15 days | 18 (7.4%) |
|  | - 16 – 20 days | 10 (4.1%) |
|  | - 21 – 25 days | 3 (1.2%) |
|  | - 26 – 30 days | 1 (0.4%) |

The cost for a patient to be treated at UKMMC A&E department for hypoglycaemia is RM 741 (USD 190). Generally, the cost of hypoglycaemic cases admitted to the medical unit is less costly than other units: RM 1,375 (USD 352) at medical unit compared toRM 1,679 (USD 430) for obstetrics and gynaecology unit, and (RM 2,611 (USD 668) atsurgical unit. Out of 903 cases with hypoglycaemia as diagnosis, 828 cases are from medical ward, 73 are from surgical ward and only 2 are from O&G unit.

As an average, the cost of care for hypoglycaemia varies with severity level. Although the mean cost of care for hypoglycaemia at severity 1 and 2 is RM7,054 (USD 1,804) and RM7,333 (USD 1,875) respectively, the mean cost reaches RM10,401 (USD 2,660) when the condition becomes severity level 3. The median cost for cases diagnosing hypoglycaemia as the primary diagnosis was RM 6,875) (USD 1,758) and cases of hypoglycaemia as secondary diagnosis was found to be RM 11,000 (USD 2,813) The median length of stay for hypoglycaemic cases was 5 days for primary diagnosis and 8 days for secondary diagnosis. In this study, the focus is on the diabetic cases with hypoglycaemia as primary reason for admission and the median cost found was RM 6,875) (USD 1,758) with a median length of hospital stay of 5 days (Table 5).

**Table 5.**
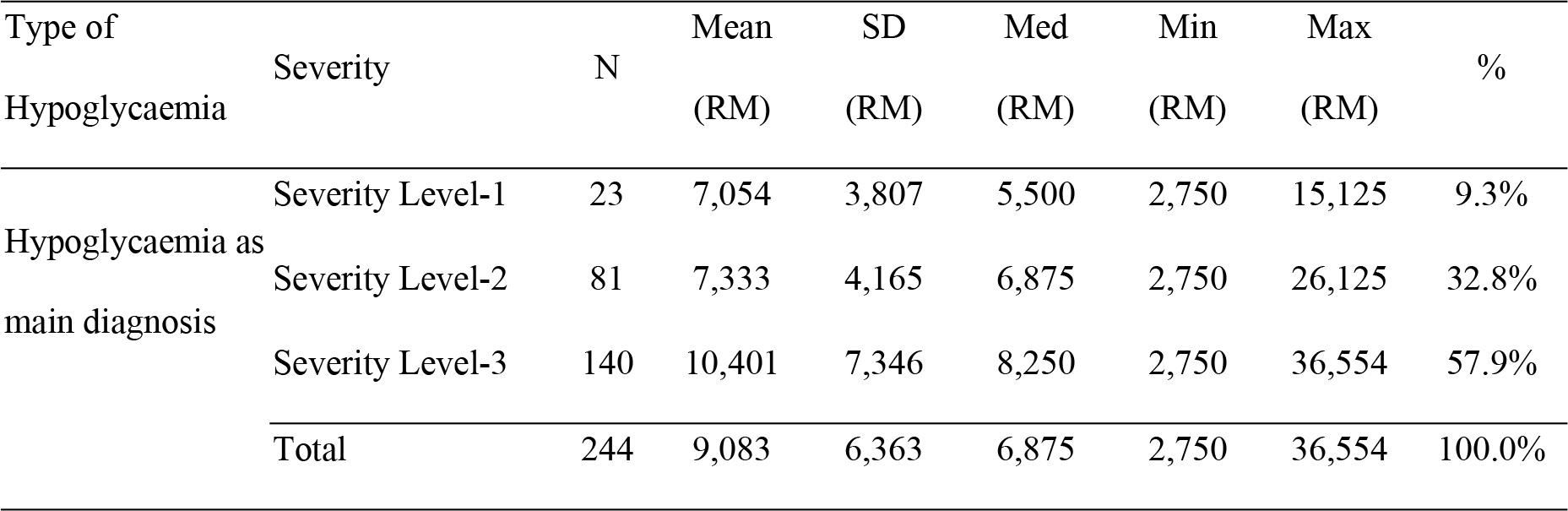
The cost (RM) of Hypoglycemia by diagnosis type and severity level

The cost by different cost centres was explored. 30% of the cost came from the final cost centre (the ward services cost), 16% of the cost came from ICU services and 15% from pharmacy and drug services.

Using the information from International Diabetes Federation [8], the HAT study [16] and the NMHS 2015 [9], the assumption on incidence and prevalence were made. From the NMHS 2015, 17.5% of adult Malaysians are estimated to have diabetes mellitus, with 8.3% of them with known diabetes. 25.1% of the known diabetics are on insulin therapy, which is equivalent to 404,619 people. Using a conservative assumption, the HAT study estimated that among type 2 DM patients, the annual incidence of any type of hypoglycaemic episode is 47.1% and severe hypoglycaemia is 16.8%. [16] Based on the assumptions on the number of cases requiring hospital admission and the mean cost for hypoglycaemia as main diagnosis in UKMMC for severity level 1 to 3, the calculation was developed as base case scenario, best case scenario and worst case-scenario. We estimated the worst case scenario based on the retrospective arm findings of the HAT Studywhich is 5.9% of severe hypoglycaemia cases (23,872 patient) assumed to require hospital admission. The base case scenario is calculated based on an expert group discussion with local clinicians, which reached a consensus of 3.2% prevalence (12,948 patients) for hypoglycaemia-related hospital admissions. In inthe best case-scenario, 2.5% prevalence (10,115 patients) was used, from the prospective arm findings of the HAT Study on hypoglycaemia hospital admissions. We assume that all these cases require hospital admission for at least once.

The total cost of care for hypoglycaemia among adult diabetes in Malaysia was estimated to be RM 117.4 (USD 30.0) million, approximately 0.5% of Ministry of Health annual budget allocation of RM 22.16 (USD 5.67) billion in 2014. [17] The national economic burden estimates based on the bestcase-scenario and the worstcase-scenario range from RM 91.7 (USD 23.5) million to RM 216.5 (USD 55.4) million. (Table 6)

**Table 6.** The Estimated Cost and Economic Burden of Hypoglycemia

| Hypoglycemia as primary diagnosis | % of cases | Mean cost per admission (RM) | Worst case scenario (5.9% admission) | Base scenario; (3.2% Admission) | Best case scenario (2.5% Admission) |
| --- | --- | --- | --- | --- | --- |
| Severity Level 1 | 9.4 | 7,054 | 15,829,227 | 8,585,371 | 6,707,321 |
| Severity Level 2 | 33.2 | 7,333 | 58,118,921 | 31,522,127 | 24,626,662 |
| Severity Level 3 | 57.4 | 10,401 | 142,522,937 | 77,300,576 | 60,391,075 |
| Total Cost |  | 9,083 | 216,471,136 | **117,408,074** | 91,725,057 |
| % MOH Budget (RM 22,160,380,300) |  |  | 1.0 | **0.5** | 0.4 |

## Discussion

Proper glycaemic control is required to minimize the risk of microvascular complication of diabetes among diabetics. At the same time, diabetes management commonly result in hypoglycaemic episodes. The occurrence of hypoglycaemia is widely variable. The frequency of severe hypoglycaemia requiring emergency services in patients with type 2 DM receiving insulin therapy is comparable to type 1 DM depending on how diabetes is being managed. Incidence rates were 11.5 and 11.8 events per 100 patient-years for type 1 and type 2 patients treated with insulin, respectively [17]. A study in Canada stated that 1.9% of individuals had at least one hypoglycaemia related A&E visits, and 0.1% was admitted to hospital. In terms of incidence rate, 5.2 cases and 0.3 cases per 1000 patient-year required A&E visits and hospital admission respectively [18]. In the HAT study, hypoglycaemic episodes of hospital admissions among DM cases in Malaysia were measured both prospectively and retrospectively. Findings from the prospective and retrospective component of the HAT study were reviewed and the most conservative estimates was selected and subsequently validated by specialists from different hospitals in Malaysia. 47.1% of the type II DM cases with insulin experienced any type of hypoglycaemia and16.8% had severe hypoglycaemic episodes within a 6-month period [16]. Based on their clinical experience, the expert group of specialists determined a 3.2% prevalence for hypoglycaemia requiring at least one hospital admission. The incidence of hospital admission was conservatively estimated as 3.2 episodes per person year and also estimated with the worst case-scenario with 10 episodes per person per year. Even with the best-case scenario, the incidence of admission due to hypoglycaemic episodes is significantly higher than elsewhere [18, 19].

Several investigations are required to confirm and manage hypoglycaemia cases appropriately. Based on severity and management protocol, the duration of hospital stay may vary from place to place. The cost of hospital care is also dependent on how long the patient was admitted to the hospital. The mean length of hospital stay ranged from 5.5 to 9.8 days in most studies (20–22). The median length of stay is 4 days in the Canada [19]. The findings from our study also provide similar estimates with the mean length of hospital stay was 6.5 days and the median length of stay was 5 days. Due to the wide variation and huge standard deviation, the median length of stay was selected as the common length of hospital stay in this study.

Depending on the level of severity, a hypoglycaemic condition may be treated at home, at A&E department or at hospital wards. The cost of care also varies with the level of severity. Our study showed that the cost of care at A&E department was RM 741 (USD 190) per case. On the other hand, the mean cost of care for a patient admitted for hypoglycaemia is RM 9,083 (USD 2,323).

The HAT study presented the proportion of patients admitted to hospital, but not the patients requiring A&E visit prior to hospital admission. Other country studies stated that the proportion of hypoglycaemia-related hospital admissions after the treatment in Emergency Department (ED) varies between 11-28% [18, 23]. We do not have local estimates for this information but based on our expert group review, it could be as high as 50%. However, we used 5.9% of the cases require hospital admission as the worst case-scenario and 2.5% as the best case-scenario from the findings in the Malaysian HAT Study [16].

Regardless of hypoglycaemia-related hospitalization is seemingly low, the actual number of hypoglycaemic episodes requiring hospital care can impose both clinical and economic burden. In this study we have focused more on the cost of care for severe episodes of hypoglycaemia at tertiary academic institutions and the total economic burden introduced by hypoglycaemia. Comparing the economic cost of hypoglycaemia is difficult due to the differences in health care systems and also in defining hypoglycaemia itself. Costing studies from the literature showed that hypoglycaemia admissions in Scotland costs USD 303 (£218) per person per day [22] whereas in Canada, where an average cost of USD 7,000 is required for an average hospitalization due to hypoglycaemia for 7 days [19]. nIn case of Thailand, a patient with hypoglycaemia requires 6 days of hospital stay as an average requiring nearly USD 700 (THB 22,000) per episode. [24] A recent study in Korea estimated the medical costs for a hypoglycaemic event ranged from USD 17.28 to USD 1,857 at secondary and tertiary hospitals [25].

Although the individual cost of care in Malaysia is not significantly high compared to other countries, the number of episodes requiring health care services at the hospital is considerably higher, making the total cost of care higher. This brings to an estimated cost of care for hypoglycaemia among type II DM patients in Malaysia to be between USD 23.5 Million to 55. 4Million (RM 91.7 Million to RM 216.5 Million). In Germany for example, the estimated annual direct cost of severe hypoglycaemia by Type II DM during 1997-2000 was USD 54,980 (€ 44,338) per 100,000 inhabitants (26,27), and in comparison, this study found the cost estimate to be significantly higher in Malaysia, amounting to at least USD 526,585 (RM 2,058,963) per 100,000 inhabitants. Although the length of stay and the unit cost of care are not necessarily higher, the number of admission required determined the possible burden for hypoglycaemic care at hospitals.

A significant portion of hypoglycaemic episodes are treated at home without the assistance of medical services either at the A&E or hospital admission [28], indicating that this study measures only the burden visible at the tip of the iceberg. Higher frequency of hypoglycaemic events could also have significant impact on quality of life as well as imposing indirect cost by limiting work capacity and work productivity.

## Conclusion

The findings of this study showed that severe hypoglycaemia in patients with diabetes impose significant impact on resource utilization. Regardless of seemingly a simple condition, hypoglycaemia can result in substantial economic burden for national health care system. Preventing hypoglycaemic episode should be included in diabetic management programs that emphasize on proper diabetic management and health education. This could minimize hypoglycaemicrisk, which may lead to reducing overall health spending, minimizing the fear of hypoglycaemia episodes and improving the compliance of diabetes management. The high cost of hypoglycaemia management calls for a personalized approach to glycaemic control and development of better guidelines for clinical decision making in diabetes control strategies.

## Acknowledgement

We would like to acknowledge the financial support we received from Novo Nordisk Pharma (Malaysia) Sdn Bhd to undertake this study. We would also like to extend our gratitude to the Research and Ethics Committee of National University of Malaysia for providing the ethical approval to conduct this research. We would also like to acknowledge Dr LetchumanRamanathan (Hospital Raja PermaisuriBainun), Prof Dr Nor AzmiKamaruddin (UniversitiKebangsaan Malaysia), Dr Zanariah Hussein (Hospital Putrajaya) for their valuable inputs in the estimated prevalence of hospital-admitted hypoglycaemia in Malaysia.

## Contributions

SM Aljunid Aniza Ismail and Yin New Aung designed the study and prepare the first draft of the manuscript. Siti Zafirah Abdul Rashid and Amrizal N Nur conducted the data analysis and prepare the tables. SM Aljunid, Julius Cheah and Priya Matzen reviewed that first draft and finalised the manuscript for publication

